# Apparent negative density-dependent dispersal in tsetse (*Glossina* spp) is an artefact of inappropriate analysis

**DOI:** 10.1101/2020.12.17.423205

**Authors:** John W. Hargrove, John Van Sickle, Glyn A. Vale, Eric R. Lucas

## Abstract

Analysis of genetic material from field-collected tsetse (*Glossina* spp) in ten study areas has been used to predict that the distance (*δ*) dispersed per generation increases as effective population densities (*D_e_*) decrease, displaying negative density dependent dispersal (NDDD). This result is an artefact arising primarily from errors in estimates of *S*, the area occupied by a subpopulation, and thereby in *D_e_*, the effective subpopulation density. The fundamental, dangerously misleading, error lies in the assumption that *S* can be estimated as the area (*Ŝ*) regarded as being covered by traps. Errors in the estimates of *δ* are magnified because variation in estimates of *S* is greater than for all other variables measured, and accounts for the greatest proportion of variation in *δ*. The errors result in anomalously high correlations between *δ* and *S*, and the appearance of NDDD, with a slope of −0.5 for the regressions of log(*δ*) on log(*e*), even in simulations where dispersal has been set as density independent. A complementary mathematical analysis confirms these findings. Improved error estimates for the crucial parameter *b*, the rate of increase in genetic distance with increasing geographic separation, suggest that three of the study areas should have been excluded because *b* is not significantly greater than zero. Errors in census population estimates result from a fundamental misunderstanding of the relationship between trap placement and expected tsetse catch. These errors are exacerbated through failure to adjust for variations in trapping intensity, trap performance, and in capture probabilities between geographical situations and between tsetse species. Claims of support in the literature for NDDD are spurious. There is no suggested explanation for how NDDD might have evolved. We reject the NDDD hypothesis and caution that the idea should not be allowed to influence policy on tsetse and trypanosomiasis control.

**Author summary:** Genetic analysis of field-sampled tsetse (*Glossina* spp) has been used to suggest that, as tsetse population densities decrease, rates of dispersal increase – displaying negative density dependent dispersal (NDDD). It is further suggested that NDDD might apply to all tsetse species and that, consequently, tsetse control operations might unleash enhanced invasion of areas cleared of tsetse, prejudicing the long-term success of control campaigns. We demonstrate that NDDD in tsetse is an artefact consequent on multiple errors of analysis and interpretation. The most serious of these errors stems from a fundamental misunderstanding of the way in which traps sample tsetse, resulting in huge errors in estimates of the areas sampled by the traps, and occupied by the subpopulations being sampled. Errors in census population estimates are made worse through failure to adjust for variations in trapping intensity, trap performance, and in capture probabilities between geographical situations, and between tsetse species. The errors result in the appearance of NDDD, even in modelling situations where rates of dispersal are expressly assumed independent of population density. We reject the NDDD hypothesis and caution that the idea should not be allowed to influence policy on tsetse and trypanosomiasis control.

## Introduction

Using critical assumptions about gene flow, and a model developed by Rousset (1), de Meeûs et al. (2) (henceforth DM) claimed that dispersal distance per generation, in tsetse flies (*Glossina* spp), increases as a power function of decreasing population density. The study was based on genetic analysis of material from five different species of tsetse, captured in ten studies in six different countries in West and East Africa (3–11). The authors concluded that negative density-dependent dispersal (NDDD) probably applied to all tsetse species. They predicted that mean dispersal distance (*δ*) would increase by 200-fold when the effective population density (*D_e_*) of adult tsetse decreased from about 24,000 to 1 per square kilometre, the order of density decline commonly associated with tsetse control operations (12). This led DM to warn that such control measures could unleash enhanced invasion of areas cleared of tsetse, so prejudicing the long-term success of tsetse control campaigns. That in turn prompted them to suggest the necessity of using “area-wide and/or sequential treatments of neighboring sites” to counter the added invasion, and they implied that small control operations risk doing more harm than good. The term “area-wide” applied to campaigns is jargon for the eradication of whole infestations up to natural barriers to reinvasion and is commonly associated with recommendations for the costly and complex use of the sterile insect technique (13). It would be dismaying if area-wide control were indeed necessary, since small-scale operations run by local communities offer an economically viable way forward (14–16).

Much therefore rests on the DM notion of NDDD and the idea thus merits prompt and thorough scrutiny. Modelling studies have already shown that, even if the NDDD hypothesis were correct, the threats to tsetse control are greatly exaggerated (17). It remains to show, however, whether there is any valid evidence that NDDD in tsetse actually exists at all. For that purpose, we now dissect the evidence adduced by DM. We show that the methods that they used to estimate their parameters are subject to massive errors, and that such errors create the illusion of NDDD, even in simulated populations where NDDD has been specifically proscribed.

## Methods

First, we identify the DM procedure and the variables that it involves. Then, employing the values that DM actually use for these variables, kindly provided by Dr Thierry de Meeûs (Supplementary File S1, Table S1), we expose anomalous relationships between them. We show that these are due to the cardinal and erroneous importance that the DM procedure gives to trap placement in the study areas, aggravated by subjective aspects of the way that such placements are viewed. By using simulations in Excel spreadsheets, we show that the errors in the DM procedure lead to the illusion of NDDD, even in simulations which specify that dispersal rates are independent of density.

The above considerations are all that need be offered to the generality of entomologists and tsetse control officers in order to invalidate completely the DM evidence for NDDD in tsetse, and thus to dispel the anxieties that DM have created. However, in the hope of interesting the specialist students of ecology, genetics and statistics, we expose many other problems in the DM arguments. We stress that we make no explicit or implicit criticism of the model of Rousset (1997), nor of the genetic analyses in any of the studies, which generated the data used by DM. We simply enumerate the errors, inconsistencies and illusions that arise from the way in which DM have used and interpreted the Rousset model and the genetic data.

## Results

### The DM procedure and its variables

DM build the idea of NDDD around their Equation (1):

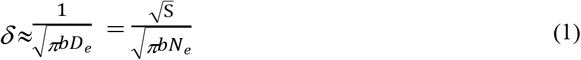

where *δ* is the predicted dispersal distance per generation of a tsetse population, *N_e_* is the effective population size, roughly defined as the number of adults in a population that will leave a genetic signature to the next generation, *S* is the surface area occupied by the effective population, and *b* is the slope of the linear regression of the genetic distance between subpopulations plotted on the log-transformed geographic distance between those subpopulations. *D_e_* is the effective population density, equal to *N_e_*/*S*. We will show that using Equation (1) in this way is inappropriate and prone to finding spurious negative correlations between the estimated values of 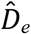 and 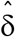. We represent estimated, as opposed to true, values of the variables by 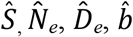, and 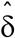.

DM did not measure 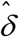: they simply predicted it from Equation (1), using estimates 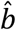 and 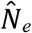 obtained from genetic analyses, and estimates 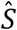 of the area occupied by the population, calculated from the disposition of the traps that sampled the population. We investigated the contributions of various factors to the value of 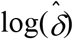 by taking base-10 logarithms of both sides of Equation (1) to get:

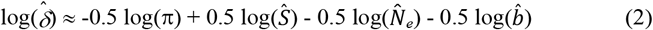

This allowed us to plot the relationships between various constituents of 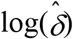 in DM’s data (Fig 1, Row A). The primary result found by DM was a strong linear correlation (*R*^2^ = 0.85; *P*<0.01) between 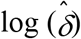 and 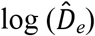 for the 10 populations used in the DM study (Fig 1, A1). DM also found a strong negative correlation (*R*^2^ = 0.86; *P*<0.02) between their predicted values of 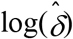 and the estimated census population densities 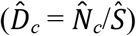, with 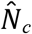 estimated from trap catches. These relationships form the central structure of the DM argument for NDDD. Notice that 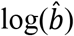 is not significantly correlated with 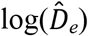 (Fig 1, A3; *P* > 0.05), while 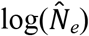 is weakly correlated with 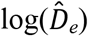 (Fig 1, A5), and not significantly correlated with 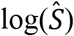 (Fig 1, A4). Surprisingly, 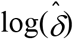 is strongly correlated with 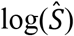 (Fig 1, A2), and 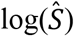 is strongly negatively correlated with 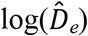 (Fig 1, A6).

**Fig 1.**
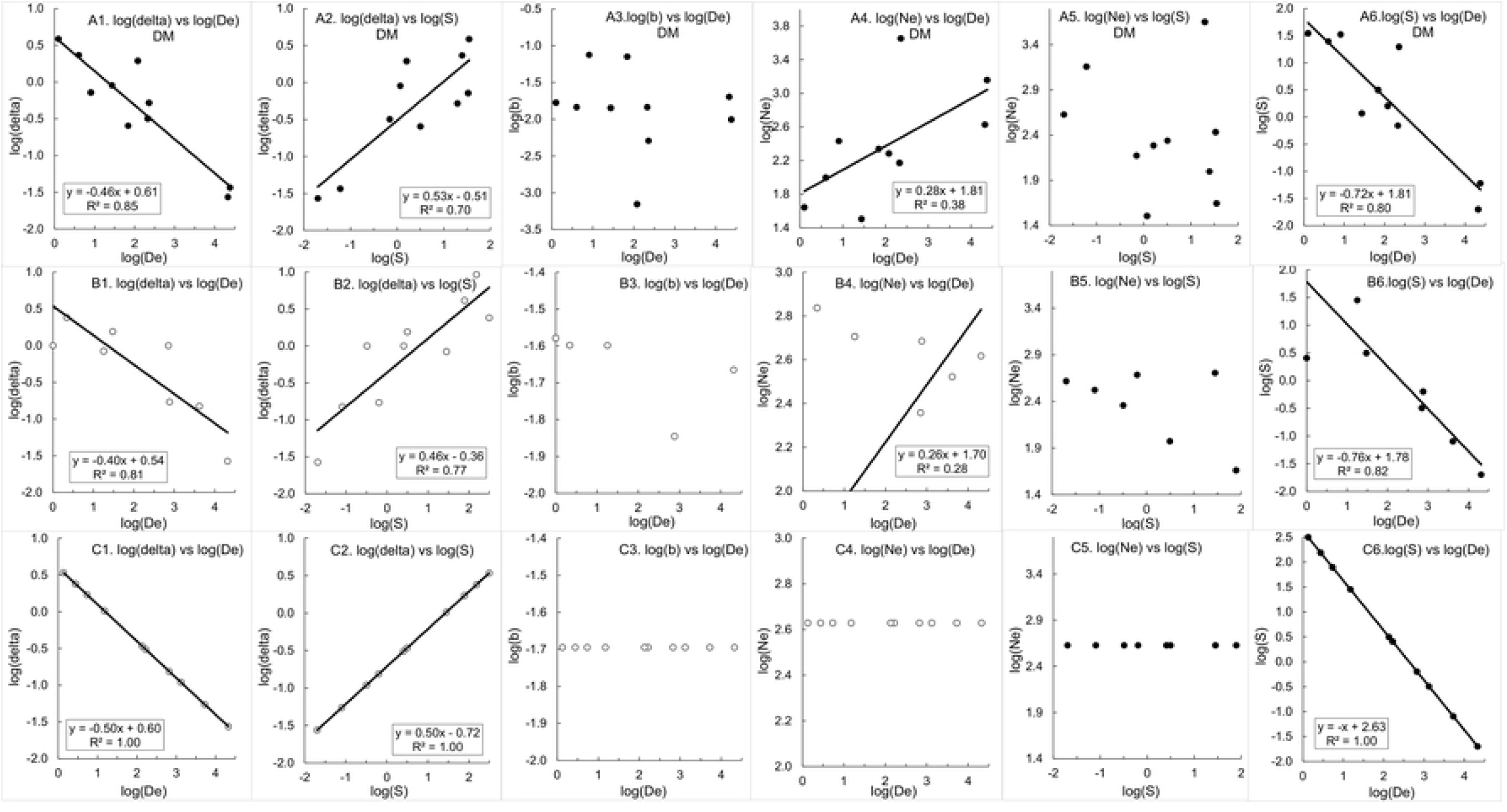
Relationships between various parameters in DM Equation (1) in real and simulated experiments. Row A: Relationships using DM estimates. Row B: Simulation using DM estimates for 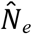 and 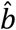 from the Opiro et al. (2017) study (11) and various values of 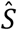 spanning the range for the 10 studies used by DM. Random error added to 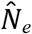 and 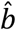. Row C: As for B but 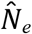 and 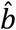 measured without error. Data kindly supplied by Dr Thierry de Meeûs and available here in Supplementary File S1, Table S1.

### Errors in the estimation of *S*, the true area occupied by a subpopulation

There is no biological reason to expect the high correlation between 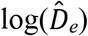 and 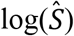 nor between 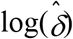 and 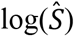. The fact that strong correlations were found, suggests that variation in 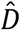 in DM’s data is mostly driven by changes in the estimated area 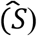 covered by traps, rather than by true changes in *D_e_*. We investigated the importance of *S* in determining the δ value predicted by Equation (2), relative to contributions from other terms. Since the equation has the form of a multiple linear regression, we borrowed a tool from regression to estimate, for the DM data, the relative importance of each of the predictor variables, 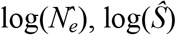, and 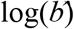, in determining the predicted value of 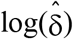.

We measure “relative importance” by the percentage of the total variance in 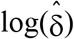 that is explained by variation in each of the three predictor variables. Since these three variables are inter-correlated across the ten DM studies, we used hierarchical partitioning (18–20) to estimate their relative importance. In this approach, one of the variables, say 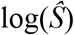, is added to each of the four possible regression models that contain neither, either, or both of the other two predictors. In each case, the increase in multiple *R*^2^ due to the addition of 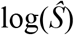 is recorded, and the average of the four increases is the estimated proportion of total variance in 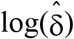 that is explained by 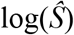. The procedure is then repeated for the other two predictors.

Because 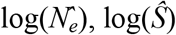, and 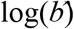 were used to predict 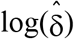 in the first place, via Equation (2), they jointly explain 100% of its total variance. Application of hierarchical partitioning to the DM data, using either the *hier.part* or *relaimpo* package of the R language, yielded the following percentages for the three predictors: 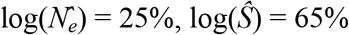, and 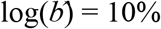. Thus, the variation in 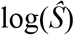 explained almost twice as much of the variation in 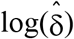 as did the other two variables combined.

The central importance of *S* in accounting for variation in the dispersal rate (δ) leads us immediately to the main problem with the DM development, embodied in their statement (2): “The average surface (*S*) occupied by a subpopulation can be computed as the surface area occupied by the different traps used in a given survey site. When only one trap was available per site or when the GPS coordinates of corresponding traps (one subsample) were not available, we computed *S = π*(*D_min_*/2)^2^ where *D_min_* is the distance between the two closest sites taken as the distance between the centers of two neighboring subpopulations”.

This statement is untrue and dangerously misleading. The true area (*S*) covered by a biological subpopulation bears no relation to the area 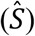 estimated to be covered by a set of traps. Assuming a subpopulation is well-mixed with random mating and no intra-population genetic structure (as is assumed by all methods for estimating *N_e_*), then *N_e_* can be estimated by catching flies from any area smaller than the geographic range of that subpopulation.

Fig 2 shows three different possibilities for the trap-sampling areas 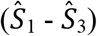 in relation to the true area (*S_t_*) occupied by the actual, well-mixed subpopulation. Sampling flies from within either 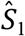 or 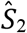 will produce the same expected value of the estimate 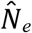, of the true value of *N_e_*, but will use different values of 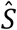, which is the denominator for calculating 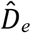. Thus, the estimated population density will vary according to the area covered by the traps, introducing error into the 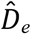 estimates.

**Fig 2.**
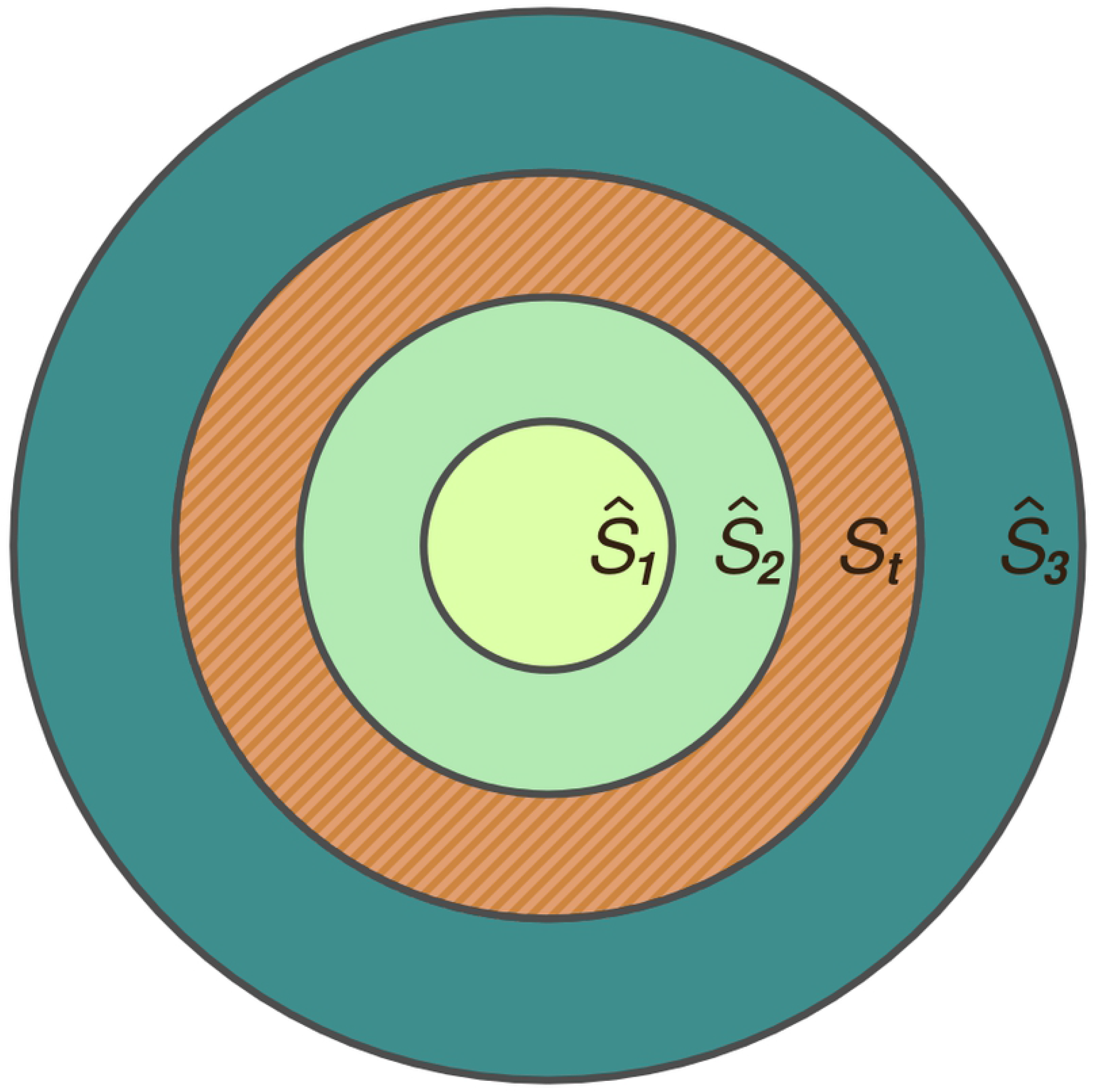
Sampled and true subpopulation areas. Illustration of three different possible sampling areas 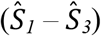 relative to the true area (*S_t_*) covered by a subpopulation.

If 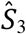 is used (covering an area larger than the size of what can be considered a well-mixed subpopulation), then the assumptions underlying the estimates of 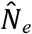 will be violated, and the estimates of 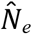 will be flawed, so introducing yet more error to 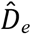. The extent to which these erroneous estimates of 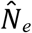 will scale with 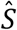 – as it grows above the size of *S* – is not entirely clear, but previous work suggests that it will not scale linearly (21) and will thus continue to produce incorrect values of 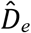 that are a function of trap placement. Correct values of 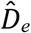 can be obtained only in the special case where *S = S_t_*. Thus, in DM’s data, variation in 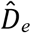 is due largely to the choice of trap placement rather than true biological variation in *N_e_*.

### How errors in 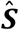 invalidate estimates of the dispersal distance 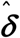

Some simple matters emerging from the DM paper allow us to check the validity of its estimates of population densities and dispersal distances. The most important of these is that, for each of the 10 populations used by DM, it is implicitly assumed that there is a single true value of δ that is roughly applicable throughout the entire area occupied by the population. Thus it should be true that, if investigators deployed traps in different patterns within the same population, but still followed the DM rules for estimating *S* in each case, the resulting estimates of δ should be the same, regardless of the trap distribution adopted – subject only to experimental errors in estimating *b* and *N_e_*. We now show that this expectation is grossly violated, even in situations where *b* and *N_e_* are measured without error.

#### In-depth analyses of the studies in Tanzania and Uganda

To see how such problems arise, consider the DM analysis of the *G. pallidipes* data for Tanzania (7). The original report states that sampling of tsetse was carried out using two traps at each of seven sites. Because GPS coordinates were apparently not available for these traps, DM analysed the data as if only one trap was used at each site. Given the definition provided above, and *D_min_* ≈ 6.7 km, they computed 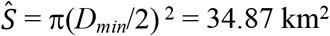. Using their estimated values of 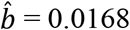 and 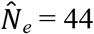, DM calculated a value of 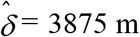 per generation from Equation (1), the highest estimate in any of the studies they cited.

As noted above, however, there were actually two traps at each site, not one. Suppose that GPS coordinates were available for the traps. Then it would be logical to use DM’s alternative definition for estimating *S*: “For all analyses, when more than one trap was available in a site, the surface area of the site was computed as *S = π*(*D_max_*)^2^ where *D_max_* is the distance between the two most distant traps in a given site, taken as the radius of the corresponding subpopulation.” If they had done that, they would have found that *D_max_* = 0.1 km, resulting in a value of 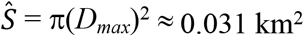, differing from the DM estimate of 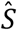 by a factor of more than 1000. Notice that, whichever definition of trap spacing was used, the same flies would have provided the genetic and geographic information employed in the DM estimates of 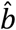 and 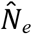, which would thus have been identical in each case. Accordingly, the revised estimates of effective population density is 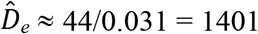 tsetse per km^2^, and dispersal distance is 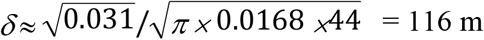, differing from the DM estimate of δ by a factor of 33.4.

This analysis suggests that the estimates DM obtained for δ, in all of their studies, were strongly dependent on the spacings of the traps used to sample the population, and also on the decision about how to interpret those spacings. Alternatively, if it is objected that – by regarding the trap deployment as either one, or two, traps per site – one is measuring the movement rates in two different subpopulations, we would be forced to conclude that values of δ could differ by orders of magnitude between subpopulations of the same population. Either scenario is sufficient to undermine completely the basis of the DM analysis.

The foregoing objections to the DM analysis do not, of themselves, explicitly negate the NDDD hypothesis as it applies to tsetse. This hypothesis is, however, seen to be similarly compromised by analysis of data from another of the studies used in the DM analysis. In a study of *G. fuscipes fuscipes* in Uganda, six traps were deployed at each of 42 sites, spread across an area of about 4000 km^2^ (11). Traps at each site were separated by a distance of at least 100 m. For this trap spacing, DM calculated 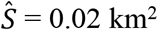, using 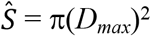 (see above). Using finite estimates of 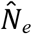, from 30 of the 42 sites, DM calculate an arithmetic mean of 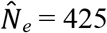 flies for the effective population (Fig 3). The other 12 sites did not provide finite estimates of 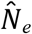. Similarly, they estimated 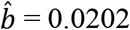 using information on genetic and geographic distances between all available sites. Using the above estimates for 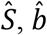, and 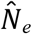, they calculated 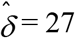 metres per generation, the lowest among all of the 10 studies cited by DM, and the one where the estimated effective population density, 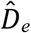, was the second highest. We now show that these estimates are also artefacts of the way in which the traps were deployed, and the way that DM chose to analyse the data.

**Fig 3.**
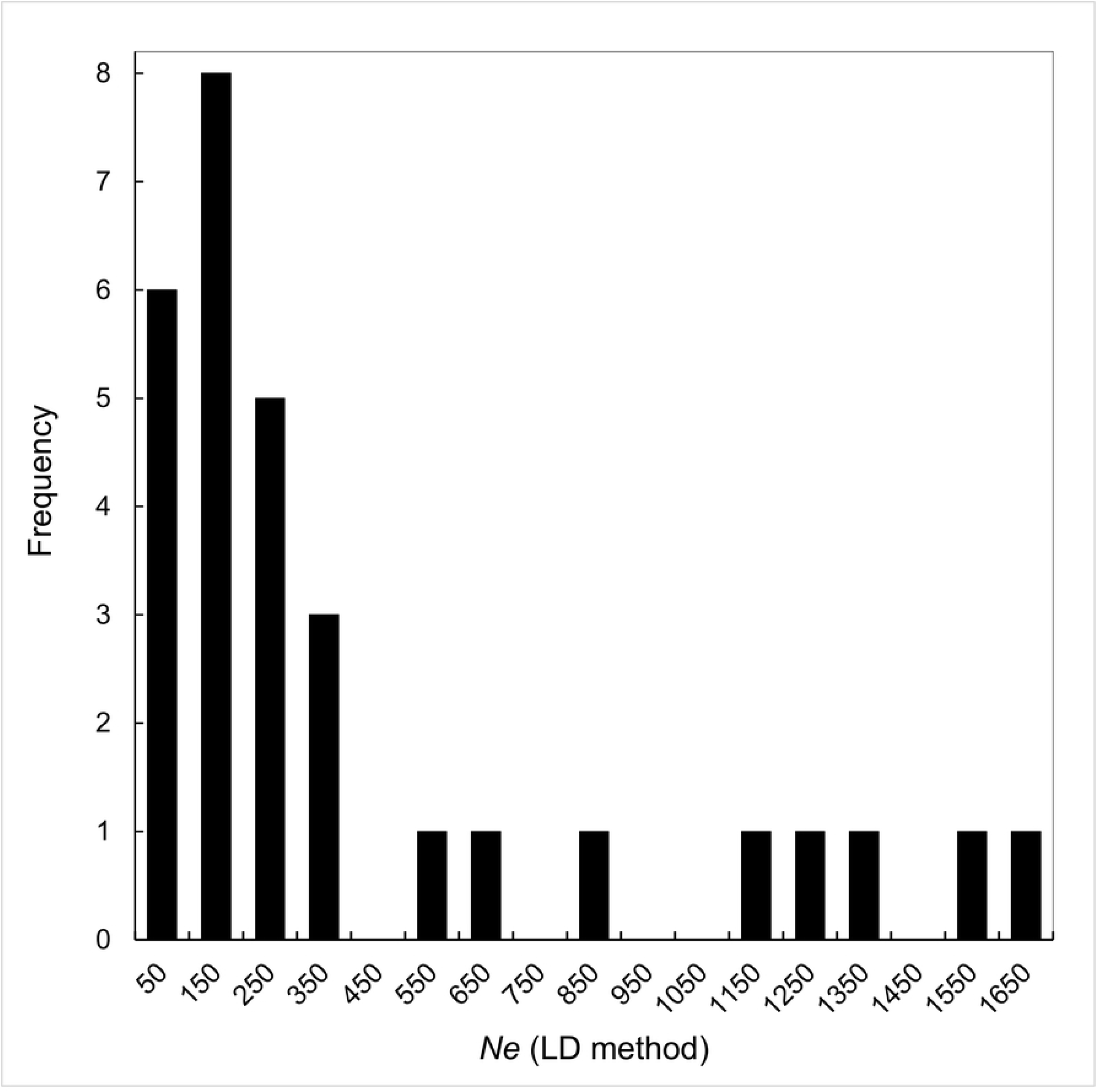
Distribution of estimates of 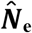 from the Opiro et al. (2017) study (11). Of the 42 study sites, 30 provided finite estimates of 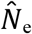, using the LD method.

To demonstrate the problem, suppose we make 10 separate estimates of the DM parameters, distributing traps throughout the roughly 4000 km^2^ in the study area (11) – with different trap deployment patterns for each of the 10 estimating procedures. For five of these procedures, suppose that – as in the original design – six traps are deployed at each site but that *r*, and thus *D_max_*, is varied such that 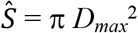 takes values of 0.02, 0.08, 0.32, 0.64 and 2.56 km^2^, respectively. For the other five procedures, larger values of 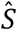 are generated by using only one trap per site, and making the shortest distance (*D_min_*) between any two sites take values of 2, 6, 10, 14 and 20 km, respectively, giving values of 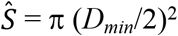 ranging from 3 to 314 km^2^. Taken together, the above design provides 10 different sampling systems – all of which are capturing flies throughout the 4000 km^2^ study area, and all of which should thus produce estimates of 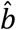 and 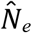 with the same expected values.

We consider two different scenarios, where *b* and *N_e_* are measured either with, or without, added stochastic error. The procedure for analysing these situations is described in Supplementary File S2W, and the calculations are carried out as in Supplementary File S2E. The results for a single realisation of the stochastic process are shown in Fig 1, Rows B and C. For each of the 10 different trap deployments, and values of 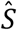, all with the same expected values 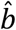 and 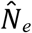, and with stochastic error added to 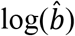 and 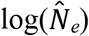, we calculate the corresponding values of 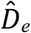 and 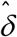.

If we assume, as DM clearly do, that the expected value of δ is constant across the whole study area (11), then the DM approach, used to analyse the data from all 10 experiments, should provide the same value of δ in each case – allowing for errors in the measurement of *b* and *N_e_*. That is to say, the measured value of δ should not depend on how the traps are deployed. In particular, if 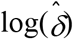 is plotted against 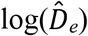 or against 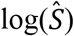, the results should approximate horizontal lines. The reality is markedly different. The results of a single realisation of the simulation procedure are shown in Fig 1, Row B, from which it is seen that the simulation essentially reflects all of the properties of the NDDD picture provided in Fig 1, Row A. In particular we see that: 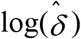 is strongly correlated with 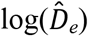, with a slope around −0.5 (Fig 1, A1, B1); 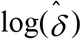is strongly correlated with 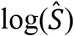, with a slope around 0.5 (Fig 1, A2, B2); 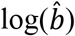 and 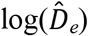 are poorly correlated (Fig 1, A3, B3); 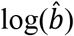 and 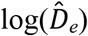 are poorly correlated (Fig 1, A4, B4); 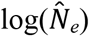 and 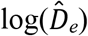 are positively, but rather weakly, correlated (Fig 1, A5, B5); 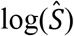 declines linearly with increasing 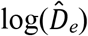 (Fig 1, A6, B6). The reader is invited to make serial iterations of the stochastic procedure – with each iteration using a different randomly generated error for 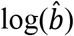 and 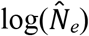.

Notice that we make no assumption about the true underlying nature of dispersal: it could be NDDD, DID (density independent dispersal) or PDDD (positive density-dependent dispersal). Regardless of this, however, the output always strongly resembles NDDD. Moreover, while we have used only one of the DM studies as an example (11), the result is entirely general and the same problem will arise in the analysis of all of the studies cited by DM. Notice also that, if *b* and *N_e_* are measured without error, the outcome still gives the appearance of NDDD (Fig 1, C1-C6). That is to say, the illusion of NDDD is entirely due to the fundamentally erroneous assumption that 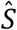, as measured by the distribution of traps used in the sampling procedure, provides a good estimate of the true area (*S*) occupied by the tsetse population under study.

The foregoing analyses show that, depending on trap placement and the choice of how 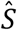 is calculated from such placement, the Opiro and Manangwa studies (7,11) can both be made to reflect either an extremely high effective population density and low dispersal rate, or completely the reverse. Clearly, in each study, these scenarios cannot both be correct. Indeed, both are almost certainly incorrect because, as explained above, 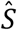 is virtually never equal to the true area (*S*) occupied by a subpopulation.

#### The illusion of NDDD in a simulated population with assumed DID

We reinforce the above results by considering the effects of errors in estimates of *S* on a simulated population where we specifically require that the true value of the dispersal distance (δ) is independent of effective population density (*D_e_*), i.e., a population exhibiting DID. Since 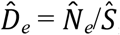, errors in the measures of *S* are reflected in the 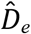 estimates and that is problematic for the DM approach. This is illustrated in Supplementary File S3, where we simulate a group of populations where δ is constant, *D_e_* varies within some arbitrary range, and *b* is calculated from δ and *D_e_* according to Equation (1). Given the assumption of DID, the calculated values of *b* decline as a power function of *D_e_* (Fig 4A). We use these true values of *D_e_* in this simulated population to generate “estimates” of 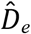 with large error. For this, 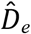 is calculated as true *D_e_* multiplied by some random factor between 0.2 and 5; these errors can be made additive instead, without consequence to the conclusion. For simplicity, we assume that 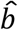 is estimated without error 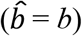. When there is error in *D_e_*, however, log(*b*) appears uncorrelated with log(*D_e_*) (Fig 4B). We then calculate 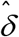 using Equation (1), replicating the method of DM. As the error input increases from 1 (no error) to 3 (three-fold error), there emerges a negative correlation between 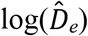 and 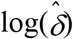, with a slope tending towards −0.5 (Fig 4C). This approximates the slope apparent in the DM analysis of their own real data.

**Fig 4.**
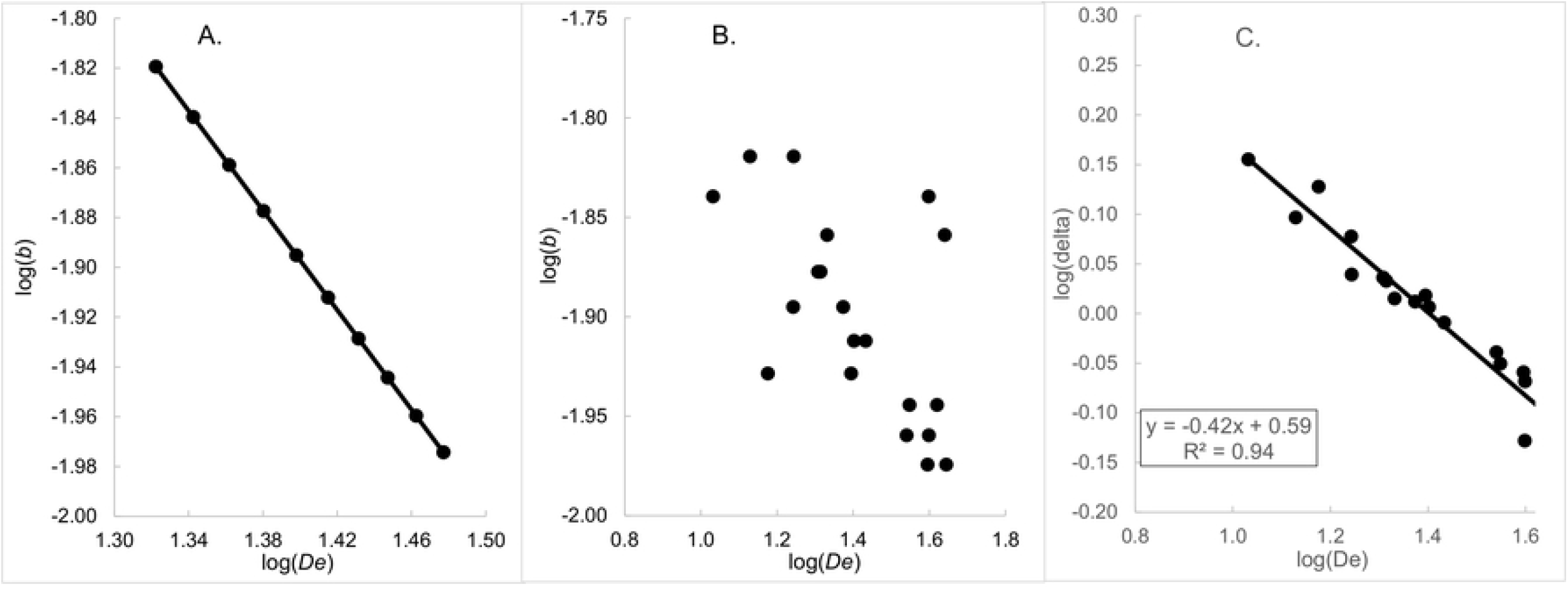
Simulation of a group of populations where the true dispersal rate (δ) is independent of the true population density (*D_e_*). A. Relationship between log(*b*) and log(*D_e_*) when *D_e_* is measured without error. B. Single realisation of the same relationship when *D_e_* is measured with error. C. Single realisation of the relationship between 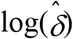 and 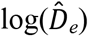 when *D_e_* is measured with error.

The reader can vary the input values of Supplementary File S3 to observe the consequences for the slope of 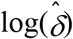 against 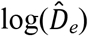. Since the simulation is stochastic, the slope changes with each realisation of the process but, for fold-error greater than around 1.2, the slope is invariably less than zero. That is to say, the population appears to exhibit NDDD, despite the fact that it has been set up such that dispersal is actually independent of population density. Furthermore, the dominating influence of large errors in 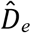 caused by highly variable and arbitrary values of 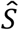, also explain the otherwise perplexing correlation we have noted between 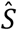 and 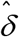 in DM’s data.

#### Why DM’s method results in apparent NDDD with a slope of −0.5

Equation (1) is a rearrangement of the original derived by Rousset (1) to describe the value of *b* that would arise in a population as a result of the values of *D_e_* and δ, where the latter is a distance measure somewhat different from that used by DM. As δ increases, the resulting genetic mixing leads to subpopulations becoming less genetically differentiated, i.e., *b* decreases, and the equality defined in Equation (1) is maintained. In the absence of density-dependent dispersal, an increase in the effective population density (*D_e_*) slows the rate of genetic drift, so that subpopulations differentiate genetically from each other more slowly, again leading to decreased genetic differentiation between subpopulations and hence a reduction in *b*. The equality defined in Equation (1) is thus maintained again, with no change in δ. If, however, dispersal is density dependent, then a change in *D_e_* will lead to a change in δ, and the resulting change in *b* will be the outcome of the changes that both *D_e_* and δ have on *b*, still maintaining the equality defined by Equation (1).

The properties above relate to the *true* values of *D_e_*, δ and *b* – and, thus, to the true values of the constituents of *D_e_* (*N_e_*, the effective population size and *S*, the area occupied by that effective population). But that is not the case for errors in the estimate 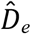. Error in 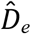 will not be accompanied by a corresponding change in 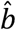, and thus, when Equation (1) is used to calculate 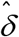 the same error will propagate to 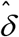, so that an overestimate of 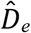 will lead to an underestimate of 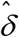, and vice-versa. When 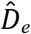 and 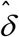 are then plotted against each-other, the result is the error in 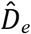 being plotted against itself. If the error in 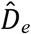 is large enough – and we have shown that errors can be >1000-fold – this autocorrelation will overwhelm any true relationship between *D_e_* and δ. As the error in 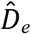 increases, 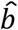 approaches independence from 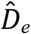, and d(ln(δ))/d(ln(*D_e_*)) can then be calculated from Equation (1) with *b* as a fixed parameter.

In Supplementary File S4 we present a complementary mathematical analysis of the effects of errors in 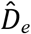 on observed results. The analysis confirms that, as long as estimation errors strongly dominate the values of 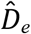, the relationship between 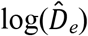 and 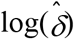 will appear to suggest the presence of NDDD, even in circumstances where the true situation is DID.

### Errors in the estimation of *D_c_*, the true census density

DM found a strong linear correlation (*R*^2^ = 0.86; *P*<0.02) between the logs of estimated dispersal distance 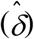 and census population density 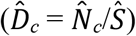, where the 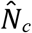 values were derived from trap catches. As we shall now demonstrate, the errors in 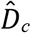 are even more serious than those in 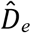. We consider a model world consisting of a continuous population of tsetse covering a large area, say 100 km ? 100 km = 10,000 km^2^. The population is uniform, across its entire extent, in terms of its true population density, which for the sake of illustration, we take as 1000 flies per km^2^. This assumption of fixed true density implies that the true effective density, *D_e_*, is also fixed, and hence δ is fixed, assuming fixed *b*. Finally, we stipulate that a single trap, used in isolation, produces a daily catch of flies equal to 1% of the population in the surrounding 1 km^2^, i.e., 10 flies per trap per day in our model world.

### Sampling the model population using more than one trap at a site

Using the DM definition for this case, *S = π*(*D_max_*)^2^, where *D_max_* is the distance between the two most distant traps in a given site. Note that this immediately assumes that the true area (*S*) of the subpopulation being sampled, and the surface area of the trapping site 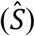, are identical. This assumption leads to problems: suppose, for example, a subpopulation is sampled using six traps deployed as shown in Fig 5A, with a single trap placed at the centre a circle of radius *r*, and a further five traps placed at equally spaced distances along the circumference of the circle. This trap placement provides the most compact disposition conforming with the trap spacing used, for example, in (11).

**Fig 5.**
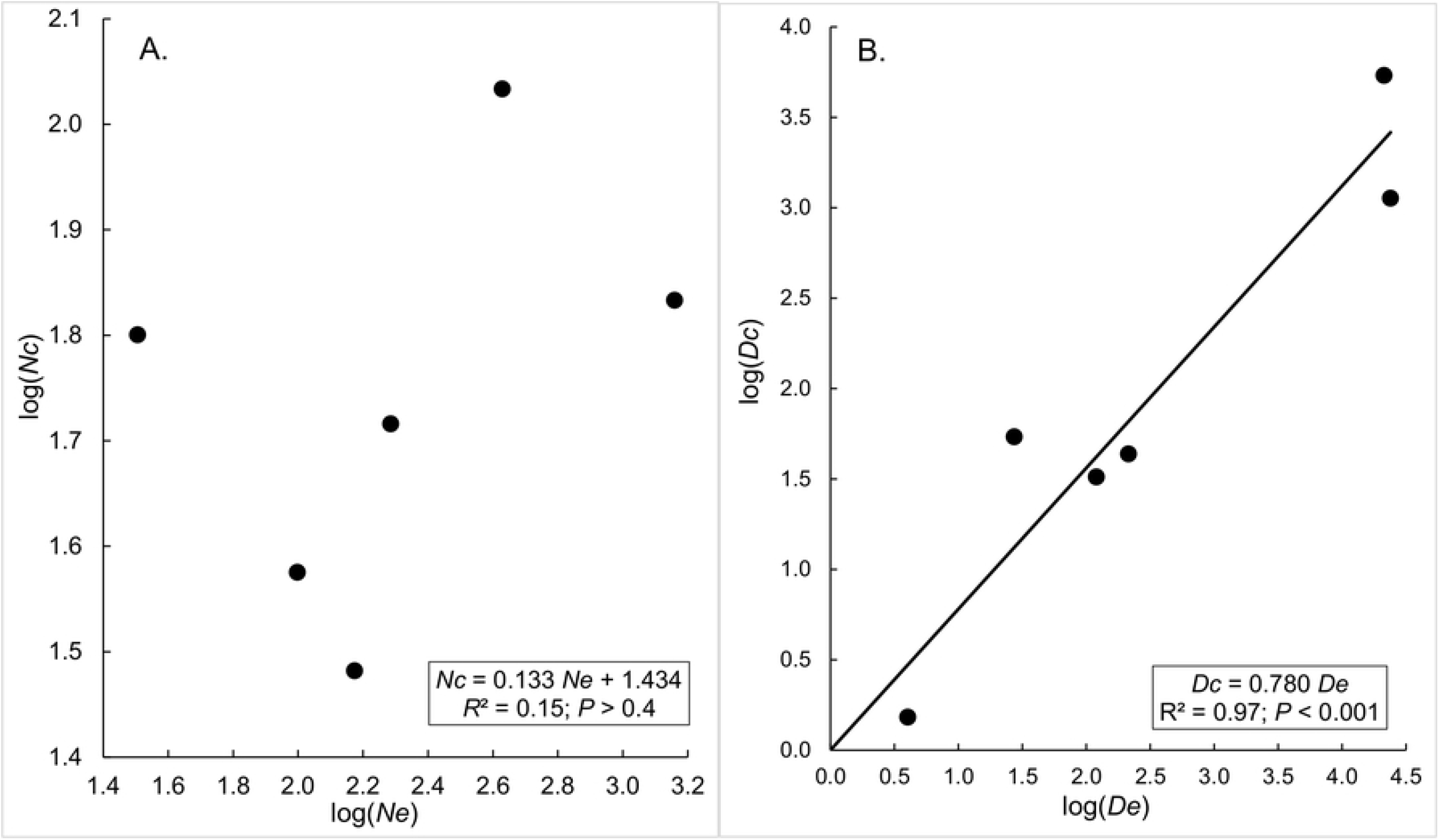
Surface area occupied by a subpopulation, as estimated by DM. A. More than one trap deployed at a site. In the example shown (*cf*(11)) five traps are spaced at equal intervals on the circumference of a circle of radius *r* units, with a further trap at the centre of the circle. DM calculate the area of the site as 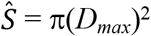, where *D_max_* is the distance between the two most distant traps in a given site, taken as the radius of the corresponding subpopulation. B. Two trapping sites with one trap deployed at each site. For this scenario, the surface area occupied by the sub-population sampled is calculated from *S = π*(*D_min_*/2)^2^, where *D_min_* is the distance the distance between the centres of two neighbouring subpopulations and thus as the average diameter of a subpopulation.

Now consider the effect of changing the value of *r*, and thus *D_max_*, on the measured values of the census population 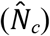 and the census population density 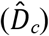 associated with the subpopulation being sampled. Initially, assume that the traps are packed very closely together, such that 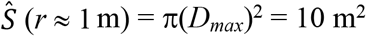. Clearly, these closely packed traps will interfere with each other and the catch per unit time would probably differ little whether one used all six traps, or just the one at the centre – which would provide an expected catch of 10 flies per day if used in isolation anywhere in our population space (see above). The expected total catch per day from all six traps would thus be *N_c_* ≈ 10 flies. If *r* is increased to 3 m, such that 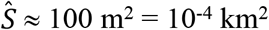, there will be less interference between the traps, but we expect that there will still be *some* interference and we thus expect 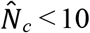 flies per trap per day, and the expected catch from all six traps will thus be 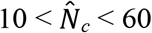 flies.

If *r* is increased to 100 m, as in (11), 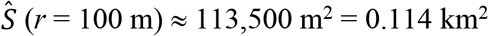, the traps are now sufficiently far apart that they are effectively acting independently (22). Each trap is, therefore, expected to catch 10 flies per day, and the expected total daily catch will be 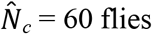 flies. For all further increases in trap spacing, and increases thereby in 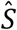, there will be no further increase in 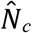 (22). Thus, the expected total daily catch is 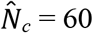 flies – regardless of the value of *r*, as long as all traps used are separated from each other by distances of the order of at least 100 m. Since, however, the census population density is given by 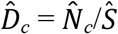 it follows that, as *r* increases above 100 m, the measured value of 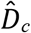 decreases as the square of the change in *r*. Thus, if we take 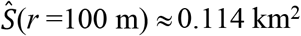 then 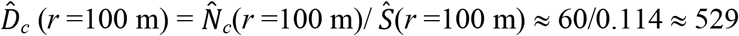 flies per km^2^. If, however, we double *r* such that 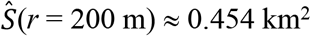 then 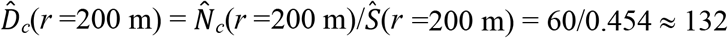 flies per km^2^, which is one quarter of the density for the situation where *r* = 100 m.

### Sampling subpopulations using one trap at each site

Using the same model world as defined above, consider a situation where we sample two sites, using one trap at each site (Fig 5B). The arguments follow the same course as for the situation where multiple traps are used at a single site. As before, the true value of δ is unaffected by the distance (*D_min_*) separating the centres of the neighbouring sub-populations. And the estimate of the census population number 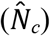 is also independent of *D_min_*, and thus of 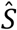. Since, however, *S* increases in proportion to the square of *D_min_*, it follows that the census population density 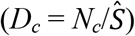 decreases in proportion to the square of *D_min_*.

That is to say, for our model world, while the catch per trap per unit time is independent of *r* ≥ 100 m, the estimated population density declines with increasing *r* – whereas the true population density is constant, being independent of the pattern of trap deployment. At the same time, however, since the catch per trap is constant, the total catch increases in direct proportion to the number of traps deployed and the duration of deployment. As evidenced by the use made of data from (11), DM took no account of these problems in their calculations of *N_c_* and *D_c_* for the various studies they used to derive the data for their Fig 1B. If this is the case, we must expect that their estimated values 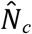 and 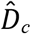, for any given study, bear little or no relation to the true values of population size and density for that study. Since, also, the various studies used to estimate 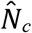 and 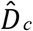 used markedly different trap numbers and spacings, it follows that the ratio of these estimated values to the true values will be different in every study. These considerations cast considerable doubt on the validity of the values of 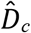 used by DM on the abscissa in their Fig 1B.

The implications of the above problems with 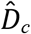 are again exemplified by contrasting the results from (11) and (7). For the former, DM use 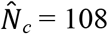, in accord with the fact that six traps, placed at least 100 m from each other, caught an average of 18 flies per trap over a period of 3-4 days, or about 5 flies per trap per day (11). Using the DM estimate of 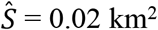 for this study, 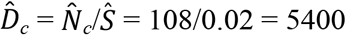 tsetse per km^2^. DM do not quote a value of 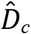 for (7) – but we know that two traps were used at each site and that each trap caught about 20 flies per day. For comparison with (11), suppose the total catch from the two traps over a 3-day period was then 2×3×20 = 120 flies. Using the DM estimate of 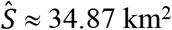 for this study gives 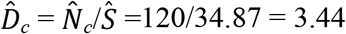 flies per km^2^. We thus have the ludicrous situation where the calculated census density is 5400/3.44 ≈ 1569 times higher in (11) than in (7), despite the catch in (11) being a quarter, i.e., 5/20, of that in (7). The apparent implication is that the availability to traps varies by at least 6000-fold between study areas, thereby suggesting that trap catches are hopeless indices of tsetse abundance. That alone is sufficient to render Fig 1B of DM entirely meaningless, although few field entomologists would credit that trap performances differ quite so markedly. The main problem, which is equally damaging, is probably erroneous estimates of 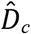 stemming from gross errors in 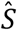.

### Absence of correlation between 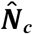 and 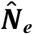

DM state that they expect a strong correlation between the true values of *N_e_* and *N_c_*. Otherwise – as they say – all population genetics studies of tsetse would need to be called into question. Their data do not, however, support their expectation. The linear correlation between their estimates of 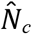 and 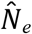 for the tsetse populations used in their own study is only 0.39 and is not statistically significant (Fig 6A). Moreover, whereas DM claim that *N_e_* values should be less than *N_c_*, in reality *N_c_ < N_e_* in five out of six situations where DM provide data for the two variables (Supplementary File S1, Table S1). Notice, however, that if we divide each 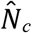 and 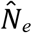 value by 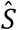, to create the population densities, 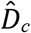 and 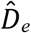, there is a very high correlation between the log transformed versions of these two variables (Fig 6B). This is consistent with our earlier suggestion that many of the DM results are artefacts resulting from inappropriate estimates of *S*, which varies over a range large enough to swamp all other sources of variation. Nonetheless, the following section suggests that there are additional good reasons to expect serious errors in the estimates of *N_c_*.

**Fig 6.**
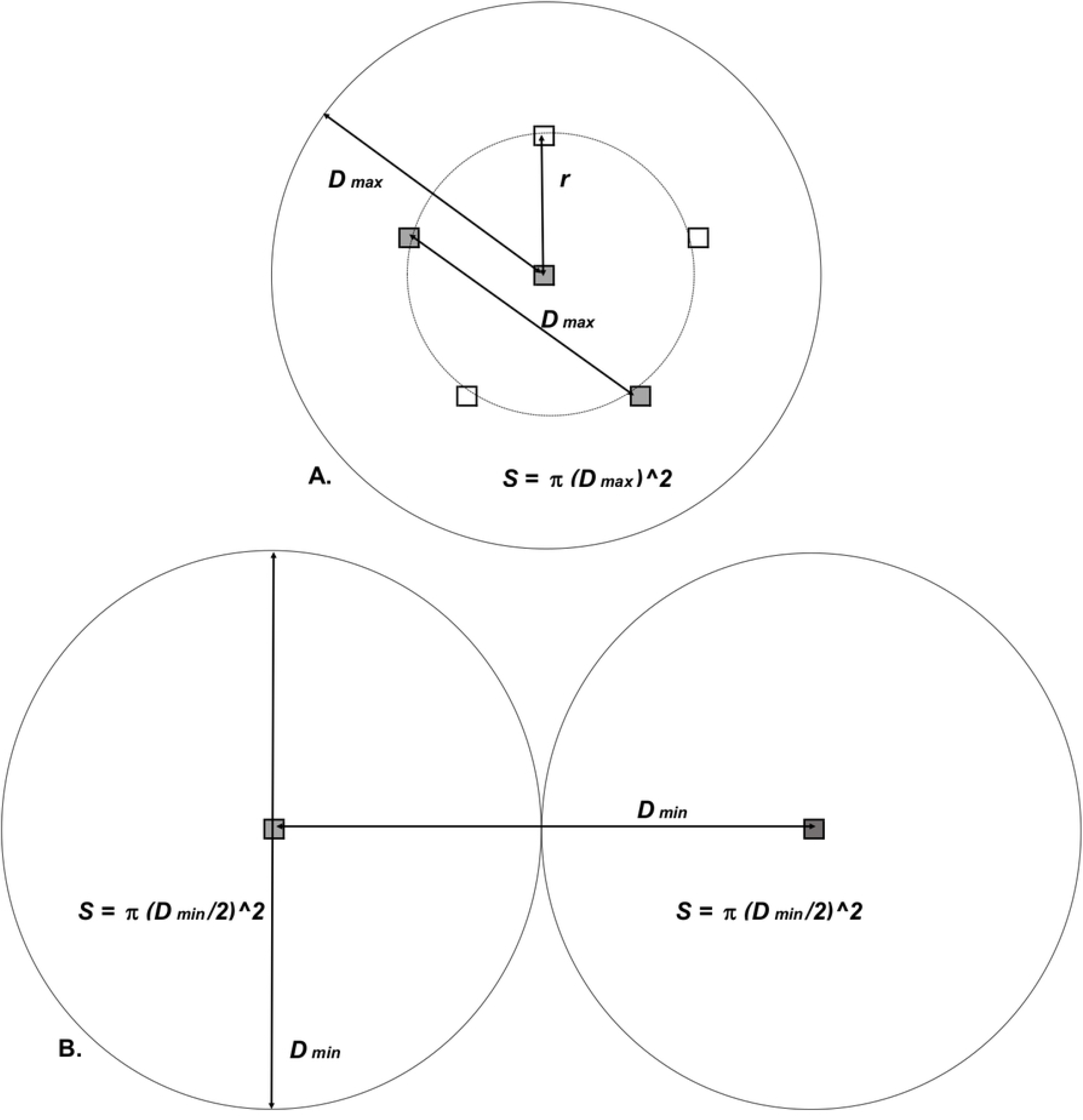
Effective and census population numbers and densities using DM data. Plots of: A. census population size 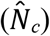 against effective population size 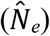; B. census population density 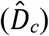 against effective population density 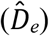. All data transformed by taking logs to the base 10.

#### Failure to allow for the intensity and duration of trapping

Scrutiny of (11) exposes further problems with the DM estimates of census numbers 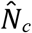 and density 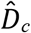. As detailed above, the DM value of 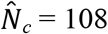 for this study reflects the number of flies captured per site – regardless of the number of traps used, and without regard to the trapping period. Failure to adjust for trapping intensity in different studies presents a serious problem. For example, with reference to Fig 5B, suppose the three traps at the white-filled squares were removed. Clearly, the expected catch – and thus 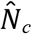 – would be halved. For the three traps remaining at the grey-filled square, however, *D_max_* – and thus 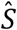 – would be unchanged, so that 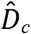 would be halved.

Moreover, there appears to have been no attempt to adjust for the trapping duration, which is approximated as an ill-defined 3-4 days. If the traps had only been run for one day, the expected catch, and 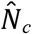, would have been reduced by around two-thirds or three-quarters. The DM paper and its Supplementary Materials provide no suggestion, for any of the study sites considered, that the catches were standardised relative to the number of traps employed and the duration of trapping.

#### Failure to allow for differences in trap performance

DM cite (23) as support for their claim that the census of flies captured in the studies they used were correlated with the real census of the corresponding tsetse populations. This is inappropriate, for two reasons. First, the paper cited provides no tsetse census data – real or otherwise. Second, it presents data only for *G. m. morsitans* and *G. pallidipes*: DM do not provide census population estimates for either species, and neither species is represented in their Fig 1B. For both of these reasons the DM census results cannot justifiably be correlated with tsetse census data from the paper cited. What the paper did suggest (23), was that there was an order of magnitude difference between the probability of capturing *G. m. morsitans* and *G. pallidipes* in an odour-baited trap (23). This underlines the danger of assuming, as DM have done implicitly in their Fig. 1B, that differences in trap catches of different species are correlated with differences between the true population densities for those species.

In offering these estimates of capture probability (23), the authors were careful to stipulate that all catches referred to a single device, run for a one-day sampling period. They also expressed the probabilities as a percentage of the population of flies occurring in a 1-km^2^ neighbourhood of the sampling device. In their own study, as detailed above, DM failed to correct their trap catches for trap type, trapping duration, numbers of traps, or the number of flies in a well-defined neighbourhood of the trap.

Even if DM had corrected the catch for the intensity and duration of trapping, there is still the problem that the studies considered by DM employed a variety of trapping techniques, involving different sorts of trap used to catch different species of tsetse. This is important because the design of traps and the species against which they are deployed can affect, by at least one order of magnitude, the numbers caught (23–27). The problem is aggravated further by the fact the some of the traps in the studies adopted by DM were employed with highly effective odour attractants (7), and others were not (11). In view of all of the above problems, it appears that the trap catches, as employed by DM, are meaningless indices of population numbers and density.

### Unsupported claims of reinvasion dynamics

When discussing possible causes of NDDD and asserting that dispersal is strongly density dependent in tsetse, DM pay no attention to the dynamics of altered dispersal that are predicted by their Equation (1). Similarly, in their abstract and on p.5, DM assert that “… control campaigns might unleash dispersal from untreated areas.” This could occur only if tsetse were reacting to the population density in the destination of their movement, i.e., the treated area, rather than in their current, untreated, location. However, this model of movement contravenes the assumptions of the underlying model (1), which clearly requires that the dispersal and density parameters must describe the same population. Thus, given that an untreated area would have a much higher population density than a treated area, flies in the former would have a lower dispersal rate than flies in the treated area. Control would not, therefore, be expected to “unleash dispersal”, that is, rapid reinvasion. DM also suggest that “… the bigger the decrease in the (treated) population, the higher potential for reinvasion…”, and this is also due to density dependence. However, this dynamic, even if correct, is not implied by their model (Equation (1)). That model says nothing about a population’s potential for reinvasion.

### Other problems

#### Errors in the estimation of *b*

Equation (1) makes sense only if *b*>0. Thus, DM tested the null hypothesis of *b=0* for each of their 10 studies, and claim that *b* is significantly greater than zero in every case. We show, in Supplementary File S5, that DM under-estimated the errors in *b* and at least three of seven estimates tested were not significantly different from zero. If all sources of uncertainty could be accounted for, it is likely that some of the other *b*-values estimated by DM would also lose their statistical significance.

#### Contradictory evidence from trap catches

The DM results often contradict common sense. An extreme example is provided by the DM estimates derived from (11), for *G. fuscipes fuscipes* in Uganda. The very low value of 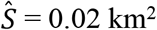, coupled with almost the highest value of 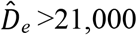 tsetse per km^2^ among all 10 studies, led to an absurdly low estimate of 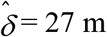 for the dispersal per generation. With a 60-day generation time assumed by DM, this is equivalent to a daily flight distance of about 4 m (17). This makes no sense: female tsetse must locate and feed on a host at least three times in a 9-day inter-larval period if they are to produce a healthy pupa (28,29). This seems virtually impossible if a fly moves only 4 m each day. The very low rate of movement is also in stark contrast to a mark-recapture estimate of 338 m/day for *G. f. fuscipes* in Uganda (30), >80 times higher than the DM estimate, and equivalent to a dispersal of δ = 1.9 to 2.6 km per generation.

Moreover, if DM are correct in claiming that census population density is at least as large as effective population density, then we might also expect that the true absolute density of tsetse in (11) could be of the order of 21,000 per km^2^. However, this makes little sense given that the results in (11) show that tsetse catches averaged only 5 per trap per day. By contrast, in (30), a single team of stationary men caught tsetse at the rate of 120 *per hour*, suggesting that the actual population density could have been 2-3 orders of magnitude higher than in (11). The problem then is that if, as suggested by the NDDD hypothesis, dispersal rates decrease markedly with increasing population density, expected dispersal rates should have been far lower in (30) than in (11), rather than the reverse.

Similar concerns are raised by the estimates derived from the study on *G. pallidipes* (7). Here the effective population density is quoted as 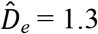 tsetse per km^2^, four orders of magnitude lower than the value estimated by DM from the study on *G. f. fuscipes* (11). It then makes no sense that tsetse catches were about four times higher in (7) than in (11). These matters all make sense, however, when one realizes that the 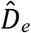 and 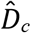 estimates in the DM study are essentially artefacts of the gross variation in trap placements and the huge resultant errors in 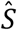.

#### PDDD more likely than NDDD

There is no credible evidence from field work with tsetse that control operations, which often cause huge reductions in population densities, result in increased dispersal rates through NDDD. In a critique of DM, an evidence-based suggestion was advanced that control operations might actually have the very opposite effect, causing what is effectively PDDD (31). Tsetse control campaigns, which raise the death rate among adult tsetse, are associated with a marked increase in the proportion of young adult females in the tsetse population (32), and such flies are unlikely to disperse far since their flight capacity is poor (33). This argument is certainly no evidence that a reduction in density will itself cause a decreased dispersal, but it does weaken further the claim that control measures will cause problems by enhancing dispersal. No effort was made to refute the above suggestion (31,34).

#### No field evidence for larviposition pheromone in tsetse

In apparent justification of the idea that females might return to previous larviposition sites, DM claim that *G. morsitans* in the field secrete a larviposition pheromone that attracts other females to the same site, leading to a strong aggregation of pupae (35). Inspection of the work cited shows, however, that it makes no mention of pheromones of any sort. Laboratory evidence for such a pheromone has indeed been published (36–38) but no such chemical has ever been shown to produce aggregation of pupae in the field. Nonetheless, the suggestion by DM that tsetse dispersal is in some way linked to the existence of a larval deposition pheromone has been reiterated by some of the DM co-authors (38). The arguments adduced in support of this idea are, however, confused or unconvincing in several ways. First, allowing that the predation of tsetse pupae increases with the local density of the pupae in the wild (39), deliberate aggregation of pupae would be expected to reduce, rather than increase, survival probability. Second, there is the potential problem that predators could benefit from a signal that pupae occur at a particular site – pupae are best hidden, not advertised. Finally, and most confusingly, it is claimed that the pheromone would have its greatest benefit at low population densities, because sparse populations become extinct if they disperse rapidly (38). This contradicts completely the DM idea that sparse populations evolve to disperse widely: in short, the DM co-authors are themselves arguing against their own notion of NDDD.

#### Inappropriate pooling of data for different situations

The DM study involved tsetse populations in ten locations, in six countries. We question the validity of performing a pooled analysis on such an *ad hoc* collection of results that refer to a variety of tsetse species in different circumstances. The pertinent question is, rather, how dispersal rates might change with the variation in population density for a given species in a given place from time to time. Hence, when DM interpret their Figure 1 as they do, they must have made the assumption that the relationship between density and dispersal rate applies to all species in all circumstances at all times. Indeed, somewhat strangely, after making this assumption to interpret their results, DM then announce that their interpretations support the assumption. However, that assumption is invalidated by our consideration (see above) of the way that the estimates of *D_e_* and *D_c_* vary due to sampling procedures and species behaviour at any given place and time. Furthermore, there are many variables other than population density that are likely to affect mobility substantially, including vegetation type and fly size (40), and perhaps also climate, and the type and abundance of hosts and sympatric species of tsetse and other biting flies.

Even if we did give credence to the procedures of DM, we would have to accept that their Figure 1 would make sense only if the habitat and population density were each homogeneous within each of the ten situations considered. Most situations in tsetse belts are not like that. For example, in the study area for *G. f. fuscipes* in Northern Uganda the terrain is stated to be very heterogeneous (11). In such conditions it is usual to place traps where experience has suggested that catches will be the greatest, ignoring the much larger areas where catches per trap are lower, but where a substantial part of the population is likely to be present at low density. Hence, to what part of the study area, if any, do the outputs for dispersal rate and density refer?

#### Absence of any suggestion for a mechanism by which NDDD might have evolved

Whether or not we allow that the population density within each study area is varied or uniform, there is a problem in seeing a credible biological mechanism for the occurrence of the NDDD apparent in Figure 1 of DM. When asked for their understanding of this, all that was offered in response was that evolution was involved, with no explanation of how such evolution was driven (31,34). However, the idea of a particular dispersal rate evolving in response to a specific density in a given situation, as conceived in DM Figure 1, creates a dilemma. On one hand, the plots of density vs dispersal in their Figure 1 would be nonsensical if the density were variable at any one time within the study area covered by each plot. On the other hand, if density were uniform within each study area at a given time, then density could not exert any direct selection pressure, since however much the flies moved, they would experience the same population density – they would not find densities more favourable to survival.

As an alternative, a combination of three phenomena might be proposed: (i) that tsetse have some optimal density, (ii) natural selection has resulted in tsetse behaving in such a way as to tend to stabilise the density at that optimum value, and (iii) the optimal density is achieved via changes in the dispersal rate, in direct response to the population density experienced by the flies. Any such change in dispersal rate would have to operate by increasing dispersal when local densities are above the optimum, and causing aggregation when local densities are sub-optimal. However, the dispersal mechanism just described is clearly PDDD, not NDDD. It is thus most confusing that, as indicated above, at least some of the DM co-authors appear eager to promote this idea (38).

#### Errors in claimed support for NDDD

DM cite various works in arguing that NDDD in tsetse populations has been known for some time (35,39,41–43). None of the papers cited, however, make any explicit or even implicit mention of NDDD; nor do they provide any support for the idea. As an example, DM cite (39) in support of the idea that the efficacy of population suppression may be reduced at low densities if normal density-dependent constraints are removed. This idea has long been accepted by the generality of tsetse workers, but it does not imply any recognition of NDDD.

#### Confusion between correlation and causation; possible reverse causality and confounding

Even if we disregard all of the many problems detailed above, and allow that the correlations apparent in Fig. 1 of DM are valid, correlation does not necessarily imply causation. Note that the gene flow theory behind Equation (1) treats dispersal and density as independent population parameters, with no suggestion of a mechanistic relationship between them (1). Moreover, even if we accepted the idea of a dependent relationship between these parameters, we could not be sure whether that meant that declining density caused increasing dispersal, or that increased dispersal caused decreased density, or that the levels of density and dispersal were each caused independently by one or more other factors. Given all of these doubts about any causal background to their Fig. 1, we conclude that DM have failed to produce evidence that tsetse control in any one place will induce any increase in the dispersal rate – let alone a gross increase. It is also presumptuous to predict that the claimed causal relationship will apply to all species of tsetse everywhere.

## Discussion

We have exposed a wide range of errors in the DM paper. Those of core importance relate to errors of measurement, which create the illusion of NDDD and ensure that the slope of 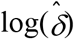 against 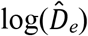 in DM’s Fig 1 approximates −0.5. DM’s method of using their Equation (1) to calculate an estimate of δ from measured estimates of *D_e_* and *b* is inherently biased towards finding NDDD (a negative correlation between *D_e_* and δ) even when none exists. This bias occurs whenever there is any error, even random, in the estimate of *D_e_*. In DM’s data, by far the most serious errors in measurement appear to be in the estimate 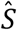, and these stem from the DM notion that the true areas inhabited by the various subpopulations under study are determined by the distance between traps. That is the root error, since inter-trap distances are not features of tsetse populations. Instead, they are strongly constrained by logistical issues, such as the numbers of available paths and traps, the mode of transport employed, the purpose of the trapping study and the whims of the researcher.

It is surprising and discouraging that no worker in the field of tsetse genetics appears to have objected to the glaring errors in the DM analysis. In particular, none of the nearly 50 co-authors on the papers cited by DM seems to have raised any concerns. We suggest that this is due to the fact that molecular genetics is a highly specialised topic, often involving abstruse theoretical considerations, arcane terminology and complex mathematical statistics, impenetrable to all but a narrow group of specialists. The consequence is that very few people have any real understanding of what is being said and implied in most mathematically-based papers on genetics. Instead, they look mainly at the implications of the results, taking on trust that the arguments and procedures which produced the results have been thoroughly checked before publication. The need for such trust is especially evident in the DM case since the procedure is particularly difficult to follow – so much so that even the editors and reviewers seem to have been overly trusting and, accordingly, misled entirely.

In consequence of all this, our willingness to credit genetic arguments in general has been weakened. We suggest that the genetics field should become more transparent, providing much clearer descriptions of analytical processes, and easy access to all of the data required to check those processes. In short, where geneticists wish to address and sway non-specialists, as DM clearly intended to do, they should be obliged to make it abundantly clear, in simple manner, exactly what they have assumed and done.

It is particularly worrying that although NDDD was initially offered by DM as a strongly supported hypothesis in need of testing, some of the co-authors are already treating the hypothesis as established fact, stating for example that: “through genetic studies, De Meeûs et al. (2019) have shown that a strong negative density-dependent dispersal occurs after control operations” (38) – clearly implying, quite falsely, that DM measured dispersal before and after control. Such statements, taken with the DM warning that NDDD will unleash massive invasion from neighbouring untreated areas, must be seen as having potentially serious impacts on disease control policy. It is for that reason that we have felt it essential to expose so fully the errors involved in the NDDD hypothesis.

Nonetheless, it would clearly be beneficial if genetic analysis could be used to provide useful indications of the dispersal rate of tsetse, or indeed of any other creature. As we have seen, a central problem is that of providing accurate and meaningful estimates of population numbers and density. That problem has taxed tsetse workers for nearly a century – since C.H.N. Jackson pioneered the use of mark-recapture in a remarkable series of population studies with tsetse (44–47) – and we still do not have good solutions. The best estimates of population numbers have involved mark-recapture exercises applied to island populations, closed to immigration and migration (48,49). When such exercises are carried out on open populations, the results are much more difficult to interpret, because of the confusing effect of fly dispersal.

Modern field studies of tsetse generally rely, as in the studies quoted by DM, on trap samples to estimate population numbers and densities, but this approach is fraught with difficulties. For example, random movement implies that a few of the tsetse caught in a trap, in a given sampling period, will have come from extreme distance, with progressively greater numbers coming from shorter distances. The area sampled in any given time is therefore a complex concept. Even if we decide to define *S* as the area from which originated some arbitrary proportion of the catch, we can assess that area only by knowing the dispersal rate of the species under study – that is, by having established in advance the very thing that procedures like those of DM, are developed to assess. It appears, therefore, that such procedures will always be difficult. Nonetheless, we are interested in collaborating with geneticists trying to solve such problems and in moving, thereby, towards a sensible use of genetic analysis in the illumination of the population dynamics of tsetse in the field.

## Acknowledgements

The contents of this publication are the sole responsibility of the authors and do not necessarily reflect the views of any funding agency. The authors declare that they have no competing interests.

